# Orthobunyavirus neurovirulence is a complex trait involving all three genome segments

**DOI:** 10.1101/2025.09.16.676581

**Authors:** Matthew J. Abbott, Christina R. Norsten, Sarah L. Anzick, Craig Martens, Clayton W. Winkler, Alyssa B. Evans

**Affiliations:** Department of Microbiology and Cell Biology, Montana State University, Bozeman, Montana, United States of America; Genomics Research Section, Research Technologies Branch; Rocky Mountain Laboratories; National Institute of Allergy and Infectious Diseases; National Institutes of Health; Hamilton, Montana, United States of America; Laboratory of Neurological Infections and Immunity, Neuroimmunology Section; Rocky Mountain Laboratories; National Institute of Allergy and Infectious Diseases; National Institutes of Health; Hamilton, Montana, Unites Staes of America

## Abstract

La Crosse orthobunyavirus (LACV) is a tri-segmented negative sense RNA virus and is the leading cause of pediatric arboviral encephalitis in the USA. The viral factors that mediate LACV’s ability to replicate and cause damage and disease in the brain (neurovirulence) are not fully understood. We previously characterized the neurovirulence of LACV and closely related Inkoo virus (INKV) and discovered they have opposing neurovirulence phenotypes in mice and human neuronal cells: LACV has high neurovirulence and INKV has low neurovirulence. We therefore generated reassortant viruses between LACV and INKV to map the genome segments that mediate LACV’s high neurovirulence phenotype. We recovered all six possible reassortant viruses of the L, M, and S genome segments using coinfection and reverse genetics approaches. We evaluated the neurovirulence of these reassortant viruses in mice *in vivo* and in human neuronal cells *in vitro*. Our results show that no single LACV genome segment alone was sufficient to cause wildtype LACV-like neurological disease in mice, and in fact all six reassortant viruses were attenuated from wildtype LACV. We found that the LACV M and S segments together were the primary drivers of neurological disease in mice, whereas the LACV L segment played a minor role. Our *in vitro* results indicate that the LACV M segment is crucial for efficient replication in neurons, but the LACV L segment appears to mediate slightly more efficient neuronal replication than the INKV L segment. The LACV M and S segments together induced wildtype LACV-like levels of neuronal death, indicating the LACV M and S are the primary mediators of neuronal death, and the L segment is not required. Together, these results indicate that LACV neurovirulence is a complex trait mediated by viral proteins on all three genome segments.

## INTRODUCTION

Orthobunyaviruses can cause severe disease in humans, including encephalitis and long-term neurological complications. The California serogroup (CSG) of orthobunyaviruses contains at least 18 mosquito-borne viruses(1). Seven of the CSG viruses have been shown to cause neurological disease in humans, including La Crosse virus (LACV) and Inkoo virus (INKV). LACV is the leading cause of pediatric arboviral encephalitis in the USA(2). INKV is primarily found in Scandinavia and has caused a small number of confirmed neuroinvasive disease cases(3, 4). The orthobunyaviruses are vector-borne and have tri-segmented, single-stranded negative-sense RNA genomes. The three genome segments are capable of reassortment between closely related viruses. Novel highly pathogenic reassortant orthobunyaviruses have been documented in nature, including Ngari virus(5–7), and the current outbreak of Oropouche orthobunyavirus (OROV) in South America is believed to be due to a novel reassortant between OROV strains(8). The potential for reassortment in dually infected mosquitoes or mammalian hosts increases with the expanding geographic range of mosquito vectors due to climate change(9). This in turn increases the chances of the emergence of either a single or reassortant orthobunyavirus. Despite this risk, it is currently unknown which LACV virus genome segments mediate the ability of the virus to replicate and cause damage and disease in the brain (neurovirulence), and how different versions of the CSG genome segments lead to different neurovirulence phenotypes. This lack of knowledge hinders the ability to develop effective treatments and vaccines(10).

The three genome segments are the L, M, and S, and together they encode six proteins. The L segment encodes the L protein, which functions as the RNA-dependent RNA polymerase (RdRp) and contains an endonuclease domain to facilitate mRNA cap snatching(11–13). The M segment encodes a polyprotein with the two envelope glycoproteins (Gn and Gc) and a nonstructural protein (NSm) of uncertain function in the mammalian host but may be involved in assembly(14–17). The S segment encodes the nucleocapsid protein (N) which forms protective ribonucleoprotein complexes with the RNA genome, and nonstructural protein (NSs)(15, 18). NSs is an interferon antagonist in many orthobunyaviruses and the only known virulence factor for LACV(19–21).

Neurovirulence depends on three primary components: 1) the ability to replicate efficiently in neurons, 2) the ability to induce neuronal death, and 3) the ability to evade or antagonize the immune system. Previous studies to determine the viral genome segments involved in neurovirulence used reassortant viruses between LACV and Tahyna CSG virus (TAHV) and mapped neurovirulence to the L segment or the M segment(22–24). However, the specific roles in neurovirulence of the genome segments were not investigated. Furthermore, these studies used highly passaged, attenuated reassortant viruses and inoculated highly susceptible suckling mice, and therefore were not a comparison of wildtype viruses and associated phenotypes in more relevant weanling or adult mouse models. A more recent study focused on the role of the LACV Gc fusion peptide in neurovirulence and the ability to gain access to the brain (neuroinvasion) by creating targeted changes to the LACV Gc fusion peptide in a LACV reverse genetics system(25, 26). These studies found that the Gc fusion peptide changes affected the ability of the virus to gain access to the CNS, but not in the ability to cause disease following direct inoculation in mouse brains and concluded the fusion protein was involved in neuroinvasion and not neurovirulence(26). Therefore, the CSG virus genome segments and proteins involved in neurovirulence and their roles are still poorly understood. Additionally, because LACV only encodes six proteins, each protein likely plays multiple roles and may be involved in both neuroinvasion and neurovirulence. Therefore, making mutations to LACV will inevitably disrupt proteins in multiple ways, making it difficult to determine the roles of LACV genome segments and proteins in neuroinvasion and neurovirulence independently.

We have previously shown that LACV and INKV share neuroinvasive ability and both are capable of crossing the blood brain barrier from the periphery to infect neurons in the brain(27). However, we have shown that LACV and INKV have differing neurovirulence phenotypes. When inoculated directly intranasally into the brains of mice, LACV causes neurological disease in all mice indicating high neurovirulence phenotype, whereas INKV causes neurological disease in few mice indicating low neurovirulence phenotype(2). Therefore, to determine the LACV genome segments that mediate neurovirulence independently from neuroinvasion, we generated and characterized reassortant viruses between LACV and INKV. The shared neuroinvasion phenotypes but differing neurovirulence phenotypes of LACV and INKV allows us to examine the genome segments involved in neurovirulence independently from neuroinvasion. We therefore generated all six reassortant viruses between LACV and INKV using coinfections and our LACV and INKV reverse genetics systems(28) and characterized the reassortant viruses in mice *in vivo* and human neuronal cells *in vitro* to determine the genome segments involved in disease, neuronal replication, and induction of neuronal death. While all six reassortant viruses were attenuated from wildtype (wt) LACV, our results indicate that the LACV M and S segments together are critical for neurovirulence, and the LACV L segment only plays a minor role in neurovirulence.

## RESULTS

### Reassortment between LACV and INKV via co-infection favors unidirectional S-segment reassortment

To generate reassortant viruses via coinfection, we co-infected Vero cells at multiple MOI ratios between LACV and INKV from 1:1 to 1:10. We initially started at 1:1 ratios, then from initial screening determined that LACV was recovered the majority of the time, so we then increased INKV in the coinfections at MOI ratios of 1:5, 1:10, and 2.5:10. For all conditions, the two viruses were co-infected at the same time on Vero cells for one hour, then cells were washed and resulting supernatants were recovered at 24 hours post infection. Supernatants were then terminally diluted, plaque isolated, then RNA isolated and screened via RNASeq for reassortment. In total, 131 plaque isolated viruses from the LACV and INKV coinfections were recovered for sequencing. Reassortant viruses are named using the first initial of the parental virus for each segment in the order of L, M, S, i.e. the reassortant virus with the L and M segment from LACV and the S segment from INKV is “LLI” virus (LLIV).

Of the 131 isolated viruses, 128 were pure cultures with a single parental origin per segment, and only three had mixtures of the same segment for both INKV and LACV, and the results are summarized in Table 1. Overall, we recovered four out of the six possible reassortant viruses: LLIV, LIIV, ILIV, and LILV. There was some variation in stock virus titers, but all four reassortant viruses grew to high titers in Vero cells indicating they were replication competent (Fig 1). Sequence summaries of the virus stocks compared to the parental viruses are summarized in Table 2. Reassortant viruses had between zero to three nucleotide changes per genome segment compared to the parental virus segments. LACV was the most frequently recovered virus and was recovered from 79 plaques (60.3%). INKV was only recovered in 11 plaques (8.4%), despite being present at higher ratios than LACV in the majority of coinfection conditions. This suggests an inherent fitness advantage of LACV over INKV, at least in Vero cells. Interestingly, the second most frequently recovered virus was the reassortant with the INKV S segment with the LACV L and M segments (LLIV), which was recovered in nearly 25% of plaque isolations. This was in stark contrast to the opposite reassortant virus, IILV, which we did not recover from any of the plaques. The remaining recovered reassortant viruses were only recovered in 4 plaques (LIIV) or one plaque (ILIV, and LILV), indicating that reassortment between LACV and INKV across most segments is not an efficient process. While we did not specifically design these coinfections to evaluate reassortment dynamics, these results suggest a unidirectional preference of the INKV S segment to reassort with the LACV L and M segments.

**Figure 1:**
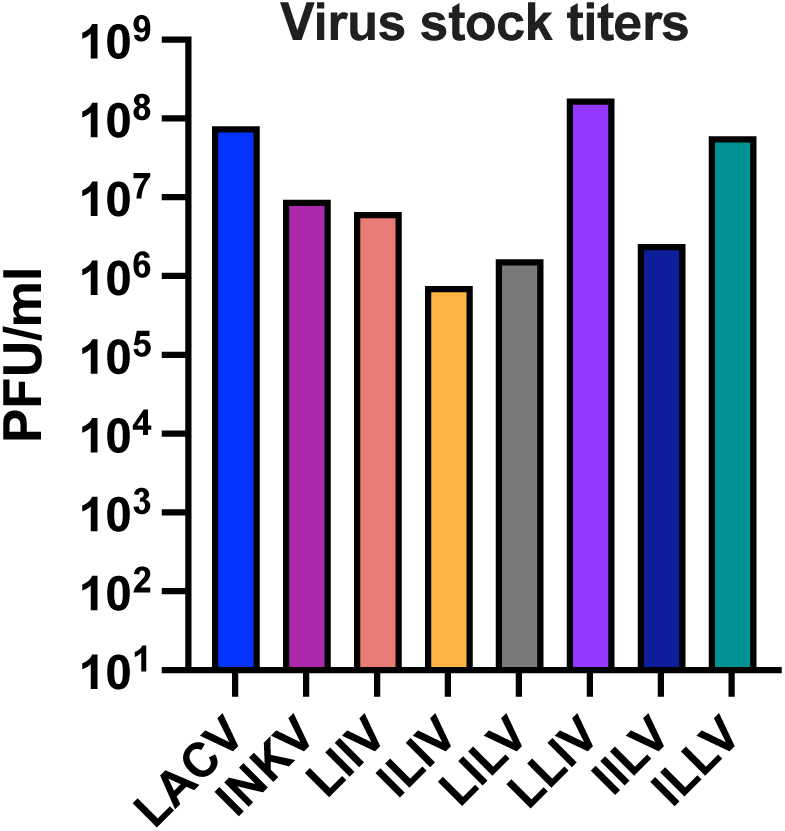
Titers of recovered reassortant and parental viruses. All stock viruses grown in Vero cells inoculated at MOI=0.1 and supernatants recovered when cells showed ≥80% CPE then titered on Vero cells.

**Table 1.**
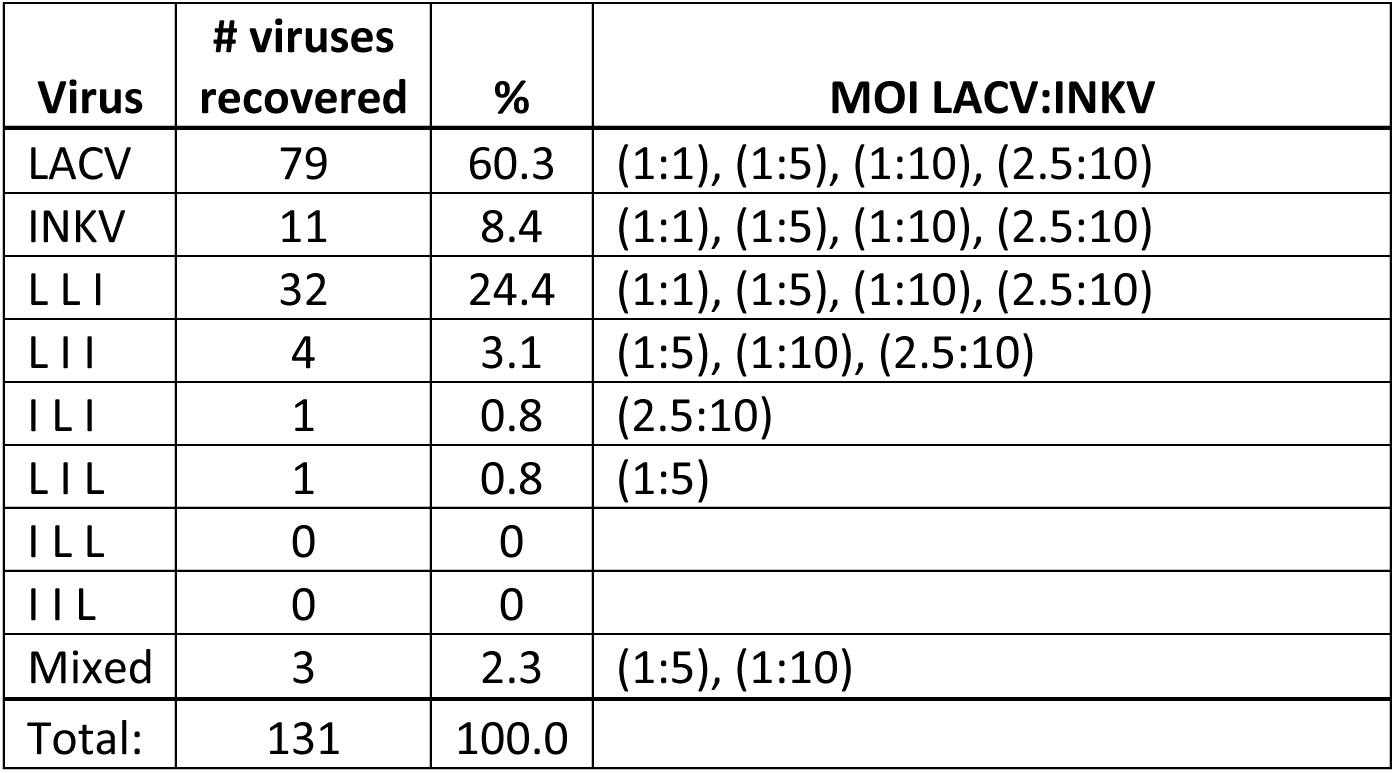
Reassortant recovery from co-infections.

**Table 2.**
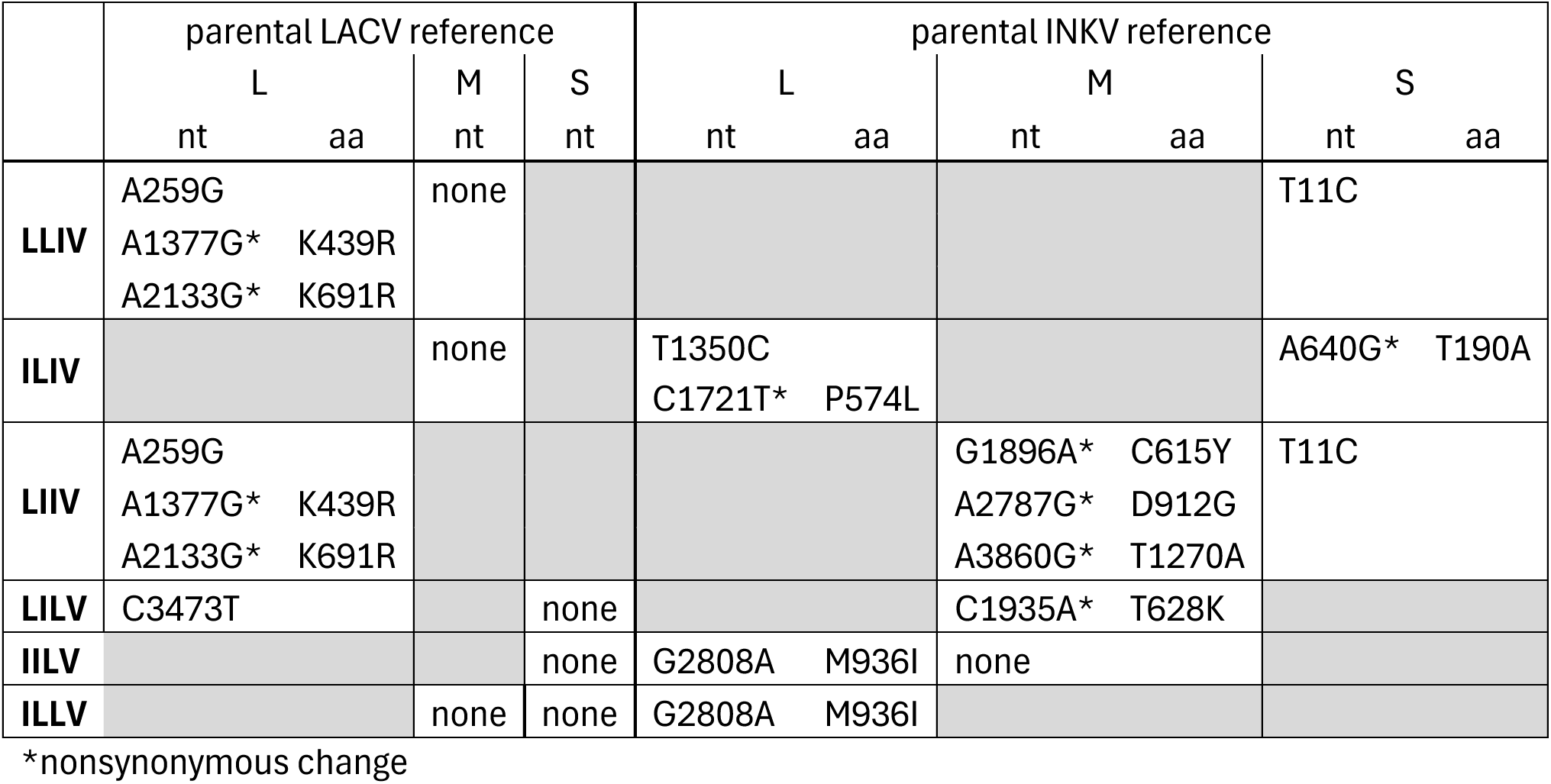
Sequence comparisons of reassortant viruses to parental LACV and INKV.

### All reassortant virus combinations between LACV and INKV are viable

The two reassortant viruses that we did not recover from the coinfections, IILV and ILLV, shared the INKV L and LACV S segments. It was unclear whether their lack of recovery was due to inefficient reassortment of these segments, or if there were incompatible interactions between the INKV L and LACV S segment that made these viruses non-viable. We therefore used our recently developed and validated LACV and INKV reverse genetics systems to generate these reassortant viruses(28). The reverse genetics systems encode a single cDNA plasmid for each genome segment for each virus under a T7 promoter, therefore through mix-and-match transfections in T7-expressing BSR-T7/5 cells, we could generate the reassortant viruses as described in(28). We successfully recovered both the IILV and ILLV reassortant viruses from the transfections and generated stocks by passage in Vero cells. Both the IILV and ILLV virus stocks had high titers in Vero cells (Fig 1). This indicates that there was not an inherent replication deficiency in viruses with the INKV L and LACV S segments together, but that they were not efficiently generated from coinfections. Sequence summaries of the virus stocks compared to the parental viruses are summarized in Table 2. Between the coinfections and the reverse genetics system derived viruses, all six reassortant viruses between LACV and INKV were recoverable and replication competent.

### The development of LACV-induced neurological disease in mice is a complex trait

We next evaluated the reassortant viruses for their ability to induce neurological disease in mice to determine the specific LACV genome segments responsible for its high neurovirulence phenotype. To do this, we inoculated adult mice intranasally (IN) with each virus at 10^4^ PFU per mouse. IN inoculation bypasses the blood brain barrier to facilitate direct entry into the brain(29), allowing investigation of neurovirulence independently from neuroinvasion. We have previously shown that LACV causes neurological disease in 100% of mice at this route and dose, whereas INKV only causes neurological disease in ∼15-20%(2, 28). Our results were consistent with our previous studies; LACV caused disease in 100% of mice at 8-9 dpi and INKV caused disease in 17% of mice at 9-10 dpi (Fig 2). Overall, all six reassortant viruses were significantly attenuated from LACV (Fig 2).

**Figure 2:**
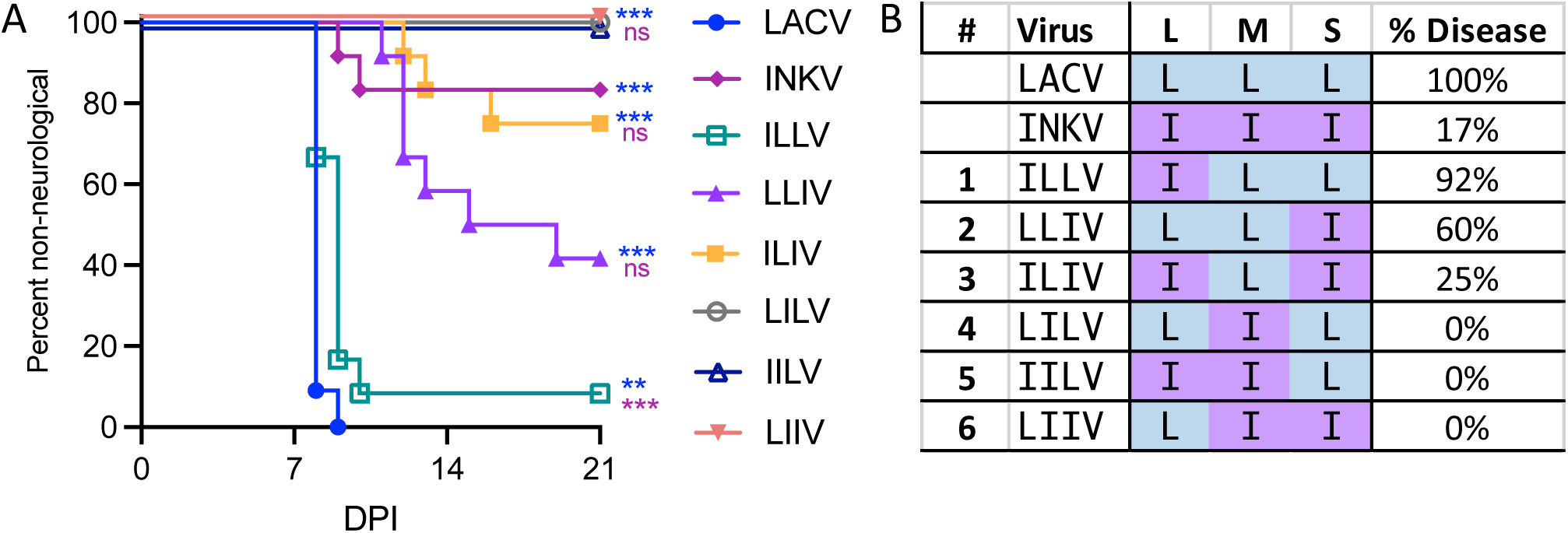
Neurological disease in mice. **A)** Neurological disease curves of adult mice inoculated IN with 10^4^ PFU/mouse. n= 11-12. *denotes results from Mantel-Cox survival test, asterisks in blue are results vs LACV, asterisks in purple are results vs INKV. **B)** Table summarizing genome segment arrangement and percent of mice that succumbed to neurological disease.

Interestingly, none of the reassortant viruses that encoded a single LACV segment with two INKV segments (ILIV, IILV, LIIV; Fig 2B #3, 5, 6) were capable of mediating wt LACV-like high levels of neurological disease in mice, similar to INKV (Fig 2A). In fact, IILV and LIIV did not cause neurological disease in any mice, and ILIV only caused neurological disease in 25% of mice. These results indicate that there is no single LACV genome segment that confers wt LACV’s high neurovirulence phenotype. In addition to IILV and LIIV, which did not cause disease in mice, the third reassortant virus without the LACV M segment, LILV (Fig 2B #4), also did not cause disease in any mice (Fig 2A). These results indicate that while the LACV M segment alone was not sufficient to cause high levels neurological disease, the LACV M segment is necessary to cause neurological disease.

The reassortant virus with LACV’s L and M together with INKV’s S (LLIV) caused neurological disease in 60% of mice (Fig 2). This was significantly attenuated from LACV, but not statistically significant from INKV (Fig 2A). However, in our extensive previous studies with INKV at this dose and route, we have never observed more than 20% of mice develop neurological disease from INKV(2, 28). Additionally, ILIV caused disease in only 25% mice, similar to INKV, and therefore the increase in disease from 25% to 60% by the addition of the LACV L segment in LLIV from ILIV suggests that the LACV L and M segments together induce more neurological disease than just the LACV M segment alone. The reassortant virus with the LACV M and S segments together with INKV’s L segment (ILLV) caused neurological disease in 11 out of 12 (92%) mice at 8-10 dpi, the most of any reassortant virus (Fig 2).

While this virus was still statistically significantly attenuated from LACV, all but one mouse developed neurological disease, indicating that the LACV M and S together facilitated a high neurovirulence phenotype similar to but mildly attenuated from wt LACV. Together, these results indicate that LACV neurovirulence is a complex trait involving all three genome segments. The LACV M and S together are the primary drivers of neurovirulence and the LACV L segment plays a minor role in neurovirulence.

### Efficient viral replication in neuronal cells is primarily mediated by the LACV M segment and aided by the LACV L segment

One of the primary components of neurovirulence is the ability to efficiently replicate to high titers in neuronal cells. We have previously shown that both LACV and INKV are restricted to replication in neurons in mouse brains, and do not infect other glial cells within the brain(2). We have also shown that LACV replicates to higher viral loads in mouse brains than INKV, even in the few INKV mice that develop neurological disease(2). Similarly, in a model of human neuronal cells, the SH-SY5Y human neuroblastoma cell line, we have shown that LACV replicates at faster rates than INKV, although INKV eventually reaches similarly high titers(2). These results demonstrate that LACV has an efficient neuronal cell replication phenotype, whereas INKV is less efficient. We therefore compared viral loads in mouse brains and replication kinetics in SH-SY5Y human neuronal cells between the parental and reassortant viruses to determine the specific LACV genome segments that mediate its efficient neuronal replication phenotype.

To evaluate replication in mouse brains, we performed time point studies in mice at pre-clinical and clinical timepoints for each virus, or corresponding time points for the viruses that did not cause much or any neurological disease. For LACV and ILLV, we took the pre-clinical time point at 6 dpi and the clinical at 9 dpi (or prior to 9 dpi if a mouse showed endpoint criteria). For the remaining viruses, we took the pre-clinical time point at 9 dpi and the second time point at 13 dpi. Overall, patterns of detectable virus in the brains followed the neurological disease curves, where viruses that caused higher rates of neurological disease in mice had higher viral loads (Fig 3). For both LACV and ILLV, which caused 100% and 92% of neurological disease in mice, respectively, high levels of infectious virus was already observed in mouse brains at the 6 dpi time point (Fig 3). There was no statistically significant difference in virus levels in the brains between LACV and ILLV at this pre-clinical timepoint. At the 9 dpi time point, 6/6 LACV inoculated mice and 5/6 ILLV inoculated mice showed neurological signs. While all of the LACV and ILLV inoculated mice had high titers of infectious virus in the brains at the 9 dpi time point, the LACV inoculated mice had significantly higher virus titers than the ILLV inoculated mice. This difference held up even if the one non-neurological ILLV inoculated mouse was removed from the analysis. This suggests that the LACV L segment, which encodes the RdRp, does help to support more efficient neuronal replication than the INKV RdRp. This slightly lower replication rate likely contributes to the mild difference in neurological disease rates in mice observed between LACV and ILLV (Fig 2).

**Figure 3.**
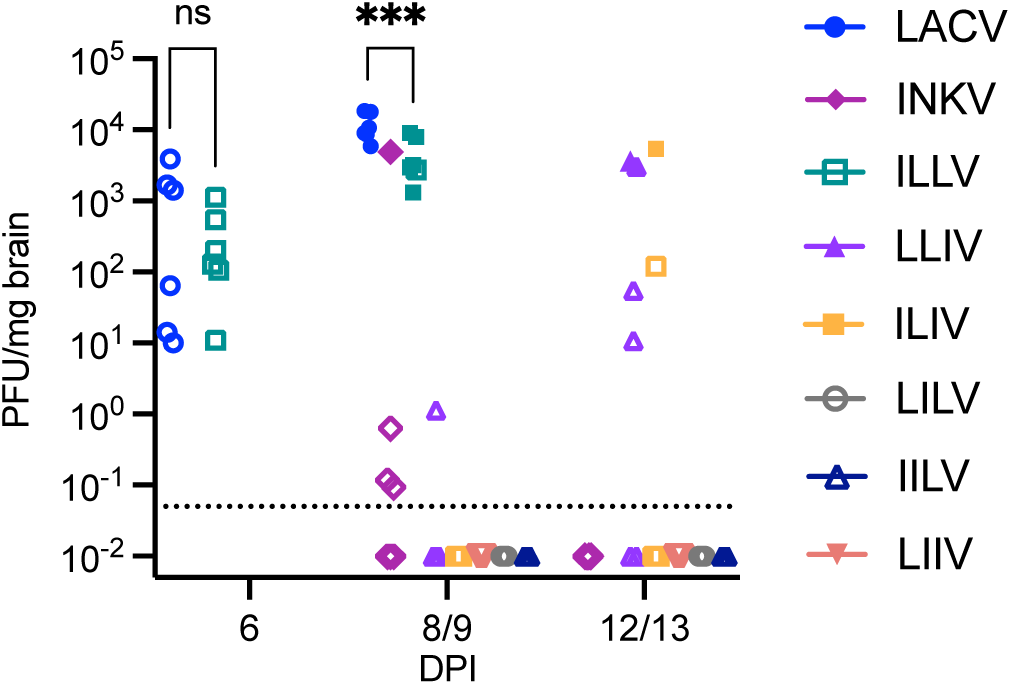
Analysis of infectious virus in the brains of mice. Time course studies of pre-clinical and clinical timepoints were conducted in mice. Brain homogenates were titered by plaque assay on Vero cells to analyze infectious virus in the brains. Viral titers between LACV and ILLV were analyzed via 2-way ANOVA in Prism. ***p-value 0.0014. Open symbols denote non-clinical mouse, closed symbol denotes neurological disease. n=6.

We next compared replication rates of the parental and reassortant viruses in the SH-SY5Y human neuronal cells. We performed replication kinetics assays by inoculating cells at an MOI of 0.01 for one hour, washing cells, replacing the media, and then collecting supernatants at 1, 6, 12, 24, 48, 72, and 96 hpi. Supernatants were titered on Vero cells via plaque assay to quantify infectious virus. Similar to our previous results with LACV and INKV, LACV replicated significantly faster than INKV, although INKV eventually reached similar end point titers (Fig 4A). While there were differences in replication kinetics, all reassortant viruses were replication competent in the human neuronal cells and replicated to similar endpoint titers as LACV (Fig 4A,B). The three reassortant viruses lacking the LACV M that did not cause any disease in mice, LILV, IILV, and LIIV, had significantly lower viral titers than LACV at 12-48 or 72 hpi, and also significantly lower rates of replication, indicating that the LACV M is important for efficient neuronal replication (Fig 4A, B). The reassortant viruses containing the LACV M segment replicated similarly to LACV, although with some significant differences. Interestingly, these differences did not fully correlate with the differences in disease rates observed in mice. LLIV only caused disease in ∼60% of mice, and yet it had the most similar replication to LACV in the human neuronal cells (Fig 4A). There was no significant difference in viral titer at any time point between LACV and LLIV by two-way ANOVA, although they did have significantly different rates of replication as analyzed via linear regression analysis of slopes (Fig 4B). ILLV, which caused 92% disease in mice, and ILIV, which caused 25% disease in mice, also replicated similarly to LACV, although they both had mildly statistically significantly lower viral titers than LACV at 48 hpi and significantly slower rates of replication from LACV (Fig 4). Analysis of LLIV, ILLV, and ILIV via two-way ANOVA showed that there is a significant difference between these three viruses (p=0.004), although there was no difference in viral titers at any timepoint via multiple comparison analysis (p>0.06). Taken together, the results from the viral loads in mouse brains and replication kinetics in SH-SY5Y cells indicate that the LACV M segment is a major contributor to LACV’s efficient neuronal replication phenotype, but the LACV L, encoding the RdRp, aids efficient neuronal replication.

**Figure 4.**
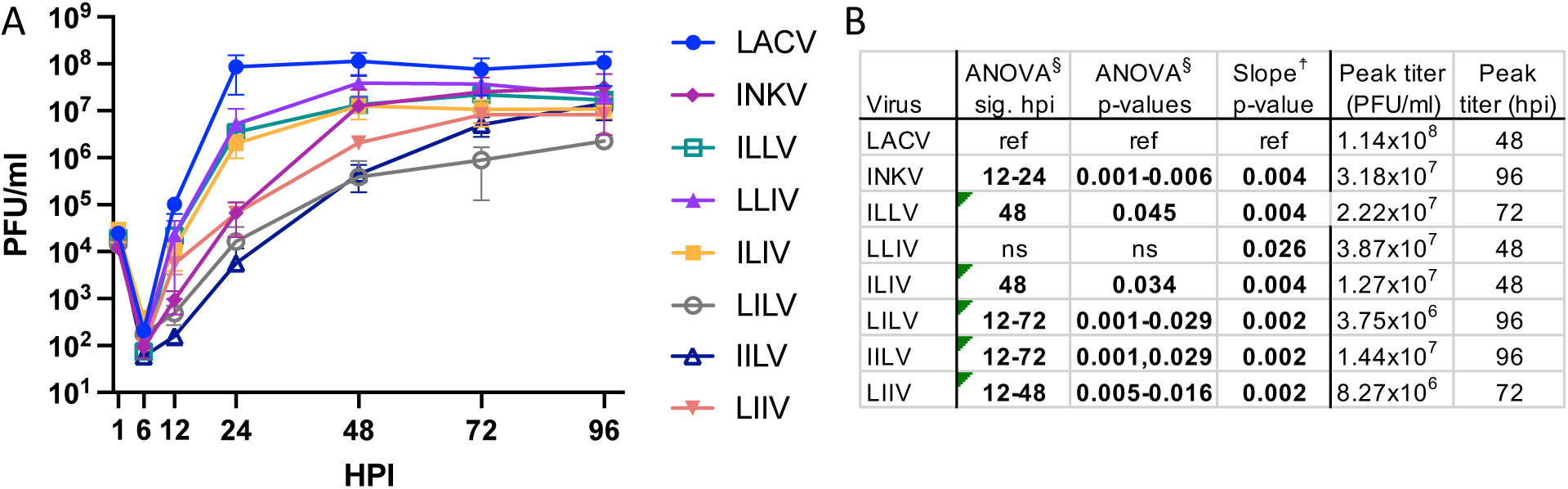
Replication kinetics in SH-SY5Y human neuronal cells at MOI=0.01. A) Cells were inoculated with virus for one hour, washed, then supernatants harvested at 1-96 hpi and titered via plaque assay on Vero cells. B) Summary of statistical analyses of the replication kinetics curves. Virus titers across time points were compared via two-way ANOVA in Prism and multiple comparisons via Dunnett’s multiple comparison test with LACV as the reference. Viral rates of replication were compared via simple linear regression analysis of slopes across the log phase of 6-48 hpi. Peak titer represents the time point with the highest average titer.

### Induction of neuronal death is mediated by the LACV M and S segments

The second important component of neurovirulence is the induction of neuronal death. High levels of viral replication may or may not induce cell death. Therefore, to examine the role of each LACV genome segment in neuronal cell death, we performed cytotoxicity assays in the SH-SY5Y human neuronal cells. We have previously shown that LACV induces high levels of cell death in these cells, whereas INKV induces little cell death(2), mirroring their neurovirulence phenotypes in mice. We again inoculated SH-SY5Y human neuroblastoma cells at MOI=0.01 of each virus and examined cell death every 24 hours at 1 to 96 hpi via CellTox green assay. The CellTox cytotoxicity assays were variable in levels of fluorescence signal from experiment to experiment, however the patterns of cell death between viruses were consistent across the four independent experiments. Similar to our previous results, INKV induced significantly lower levels of cell death than LACV by 96 hpi (Fig 5). LACV started to induce cell death by 48 hpi and induced very high levels of cell death by 96 hpi (Fig 5). The reassortant viruses without the LACV M segment that caused no disease in mice and had slow replication phenotypes in the SH-SY5Y cells (LILV, IILV, and LIIV) induced significantly lower levels of cell death than LACV, similar to INKV, through 96 hpi (Fig 5). Interestingly, while the reassortant viruses ILLV, LLIV, and ILIV had similar replication phenotypes in SH-SY5Y cells, there were distinct differences in their ability to induce death in SH-SY5Y cells. While the two-way ANOVA did not identify any significantly different time points between LACV and ILLV, LLIV, and ILIV, likely due to the variability in fluorescence across experiments, we observed consistent patterns in cell death across the four experiments that are represented in the combined graph (Fig 5). The reassortant viruses with the LACV M alone (ILIV) and the LACV L and M together (LLIV) induced intermediate cell death that was consistently higher than the INKV levels but lower than the LACV levels of cell death (Fig 5A). Interestingly, the virus with the LACV M and S together, ILLV, consistently induced high levels of cell death, nearly identical to LACV (Fig 5A). Together, these results indicate that the LACV M and S together mediate high levels of neuronal death, and the LACV L likely does not play a large role in cell death.

**Figure 5:**
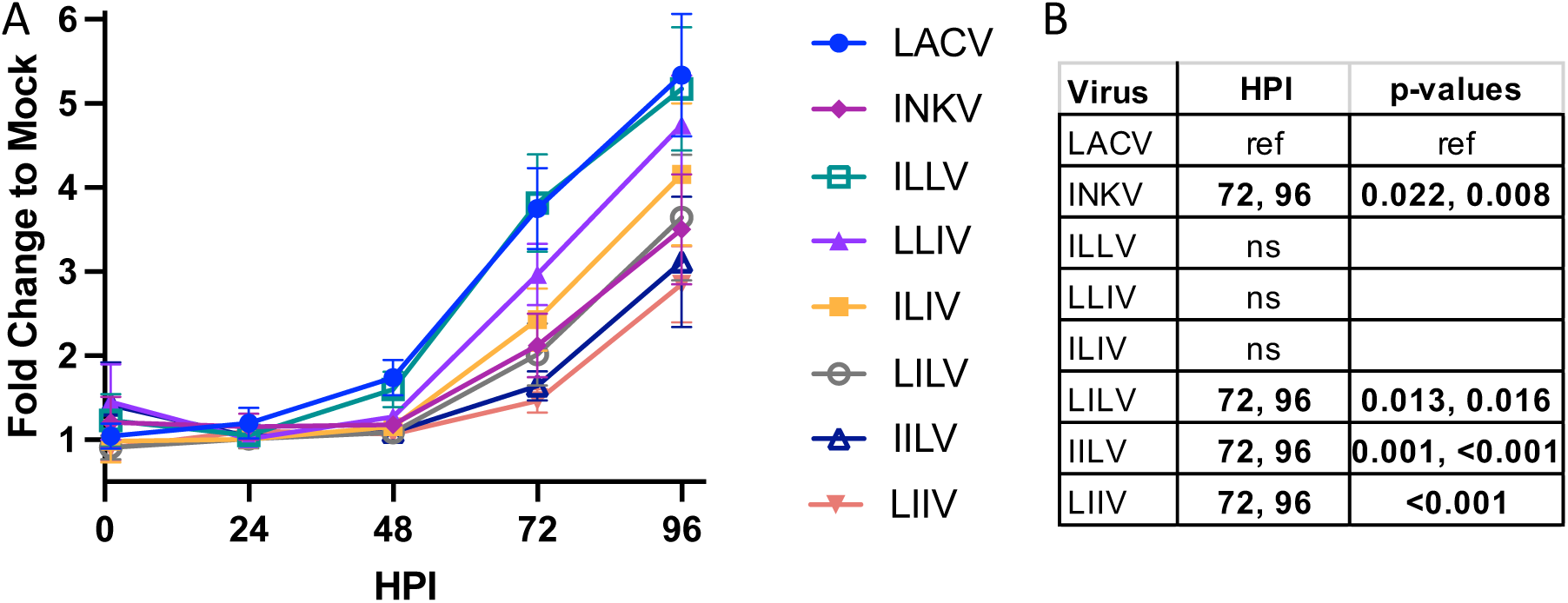
Cytotoxicity assays of LACV, INKV, and reassortant viruses in SH-SY5Y cells at MOI=0.01. Cell death analyzed via CellTox Green Assay (Promega). A) Fluorescence graphed as fold change to mock-inoculated control wells with standard error of the mean. B) Summary of two-way ANOVA comparing cell death across time with LACV as the reference. Results represent four independent experiments.

## DISCUSSION

Leveraging reassortant viruses between two CSG viruses with different neurovirulence phenotypes allowed us to examine the orthobunyavirus genome segments that mediate differences in neurovirulence. Neurovirulence depends on three separate components: 1) the ability to replicate efficiently in neurons, 2) the ability to induce neuronal death, and 3) the ability to evade or antagonize host immune responses. In this study, we examined reassortant viruses for their ability to cause neurological disease, as well as their role in the individual components of efficient viral replication and induction of cell death in neuronal cells. We have shown that all three LACV genome segments are involved in LACV’s high neurovirulence phenotype, but the segments appear to contribute in different ways.

Overall, the LACV M segment was critical for neurovirulence, however alone it was not sufficient for inducing neurological disease in mice, efficient replication in neuronal cells, or inducing cell death as evaluated by the ILIV reassortant virus (Figs 2, 4, and 5). These results are consistent with a previous study from the 1980s that found M to be a major mediator of LACV neurovirulence in suckling mice(24). Our results showed that the LACV M segment was the most important LACV segment for efficient replication in neurons, as the viruses lacking the LACV M (INKV, LILV, IILV, and LIIV) all had significantly reduced viral titers across multiple time points and replication rates compared to LACV, although they all reached similarly high titers by 96 hpi (Fig 4A, B). Perhaps unsurprisingly given their lower replication rates in neurons than LACV, these viruses also induced low levels of cell death (Fig 5), indicating that the LACV M segment is also crucial for neuronal death. The reassortant viruses containing the LACV M segment (ILLV, LLIV, and ILIV) replicated more similarly to LACV in human neuronal cells, with only ILIV and ILLV differing in viral titers from LACV at the 48 hpi time point (Fig 4B). Interestingly, despite the similar neuronal replication phenotypes of ILLV, LLIV, and ILIV, these viruses differed substantially in their ability to induce neuronal death. They all induced higher levels of cell death than INKV, but the LACV M segment alone (ILIV) did not induce the same high levels of cell death as LACV. This indicates that the LACV M segment is critical for the induction of neuronal death, but is not sufficient on its own to induce the high levels of neuronal death and requires interactions with other viral components. The results from mice and human cells indicate that the LACV M segment is critical for the induction of neurological disease in mice by facilitating both efficient replication in neurons and high levels of neuronal death. Further research is needed to determine the specific roles of Gc, Gn, and NSm.

The LACV S segment was critical for high levels of neurological disease in mice and induction of high levels of cell death (**Figs 2, 5**), however it was unnecessary for efficient replication in neurons (Fig 4). The LACV S segment alone (IILV) was insufficient for causing neurological disease in mice, efficient neuronal replication or high levels of neuronal death, and only exhibited high neurovirulence in the presence of the LACV M segment (ILLV). This indicates that there are interactions between proteins on the LACV M segment and proteins on the LACV S segment that facilitate neuronal death and high levels of disease in mice, either through direct protein-protein interactions or complementation of function. Previous studies on Bunyamwera orthobunyavirus (BUNV) have shown that the BUNV N protein encoded by S directly interacts with the glycoproteins encoded by M, however the nature and function of this interaction is unknown(17). Perhaps these interactions help trigger neuronal death, although further research into these interactions is required to determine their role. Additionally, the S segment encodes NSs, which is an interferon antagonist in LACV(20). It is possible that the NSs could play an additional role in cell death. Alternatively, the LACV NSs may better antagonize the interferon response in mice than the INKV NSs, and this could lead to the higher levels of neurological disease observed in mice with the LACV M and S (ILLV) than just the LACV M (ILIV; Fig 2). Our results clearly show that the LACV S segment is an important driving factor for neurological disease in mice and high levels of neuronal death, but future work on N and NSs is required to determine their mechanistic roles.

The LACV L segment, which encodes the L protein including the RdRp, played the least critical role in neurovirulence. However, the LACV L was required for wt LACV’s full neurovirulence phenotype in mice (Fig 2). The virus with the LACV L and M together (LLIV) induced neurological disease in 60% of mice versus in only 25% of mice with the virus without the LACV L but with the LACV M (ILIV). Similarly, the virus lacking only the LACV L (ILLV) was highly neurovirulent in mice (92% neurological disease) but was still mildly attenuated from LACV (100% disease), together suggesting that the LACV L does contribute to LACV’s neurovirulence. Additionally, evaluation of viral loads in mouse brains from mice inoculated with LACV and ILLV displaying neurological signs showed that there were significantly lower viral loads in the ILLV mice than the LACV mice, indicating there was slightly less replication efficiency without the LACV version of the RdRp (Fig 3). Similarly, in the SH-SY5Y human neuronal cells, the LLIV reassortant virus was the only virus with indistinguishable titers from LACV across all time points (Fig 4B), whereas the virus without the LACV L but with LACV M (ILIV) had significantly lower viral titers than LACV at 48 hpi. However, LLIV and ILIV did not have significantly different viral titers from each other at any time point. Interestingly, despite the slightly more efficient replication of LLIV compared to ILIV, these viruses induced very similar levels of neuronal death. The virus with the most similar cell death to LACV was ILLV, which lacks the LACV L. This indicates the LACV L protein is not critical for cell death. Taken together, the results from mice and human neuronal cells suggests that the LACV RdRp contributes to slightly more efficient virus replication in neuronal cells compared to the INKV RdRp, which results in only a mild difference in neurovirulence phenotype.

Together, our results indicate that LACV neurovirulence is a complex trait requiring all three genome segments. The LACV M segment facilitates efficient neuronal replication which is mildly aided by the LACV L segment, and the LACV M and S segments together facilitate high levels of neuronal death which is correlated with high levels of neurological disease in mice. Future work is required to determine the specific viral proteins that mediate these processes and the mechanisms involved.

## METHODS

### Cells, viruses, and plasmids

Vero cells (ATCC) were maintained in DMEM (Gibco) supplemented with 10% FBS (Atlas Biologicals) and 1% penicillin-streptomycin (Gibco). SH-SY5Y cells (ATCC) were maintained in 1:1 ratio of EMEM:F-12K (ATCC) supplemented with 10% FBS and 1% penicillin-streptomycin. BSR-T7/5 cells were maintained in GMEM (Gibco) supplemented with 2mM L-Glutamine solution (Gibco), 1x MEM amino acid solution (Gibco), and 10% FBS. BSR-T7/5 cells were passaged under Geneticin (Gibco) selection at 1mg/ml every other passage.

Wild type parental virus stocks grown in Vero cells of LACV (human 1978) and INKV (SW AR 83-161) were used and previously described in(2). The LACV and INKV reverse genetics systems were previously described in(28). All reassortant viruses, whether generated from coinfections or the reverse genetics system, were passaged on Veros in a six-well virus mini stock in 2.5ml DMEM supplemented with 2% FBS, 1% pen-strep, and 25μg/ml of Plasmocin (Invivogen) and harvested at ≥80% CPE. Final stock viruses were grown in Vero cells at inoculation of MOI=0.1 for one hour, cells washed, then fresh DMEM + 2% FBS + 1% pen-strep (“virus media”) added. Supernatants were harvested at ≥90% CPE, clarified at 500xg for 7 minutes, and aliquoted. Clarified virus stocks stored at −80°C.

### Virus titers

All viruses were titered in Vero cells via plaque assays. Viruses were serially diluted in virus media and 200μl plated per well in duplicate onto 24-well plates of Vero cells plated the previous day at 1.3×10^5^ cells per well. Plates were incubated for one hour at 37°C and then overlayed with MEM (Gibco) with 1.5% carboxymethyl cellulose (CMC; ThermoSci) and incubated at 37°C. At 4dpi, cells were fixed with 10% formaldehyde and stained with 0.35% crystal violet and plaques counted.

Viral titers from mouse brains were performed similarly using brain homogenates. Brain homogenates were prepared using half brains cut sagittally. The half brains were weighed and homogenized in 2 ml cryovials (Sarstedt) containing 2.3 mm Zirconia/Silica beads (Fisher) in 0.5ml serum-free DMEM. Bead homogenization was performed on a bead beater at 5300 rpm for 25 seconds, then the samples were clarified at 5000xg for 10 minutes at 4°C, and the clarified supernatant used for virus titering on Vero cells as described above.

### Generation of reassortant viruses

This work was approved under the NIH Rocky Mountain Laboratories IBC registration 24-RML-006. The project and manuscript were reviewed and approved by the NIH DURC-IRE. This work was additionally approved under Montana State University Biosafety protocol 2023-444-IBC. The reassortant viruses LLIV, LILV, ILIV, and LIIV were recovered from coinfections of LACV and INKV in Vero cells (ATCC). For coinfection reassortant recovery, Vero cells were seeded at 5×10^5^ cells per well in a 6-well plate the previous day. Cells were co-infected with LACV and INKV at ratios of 1:1, 1:5, 1:10, and 2.5:10, respectively. Cells were incubated with the virus for one hour, cells washed with media, then replenished with 2.5 ml virus media. Supernatants were harvested and clarified at 24 hours post inoculation (hpi). For initial terminal plaque dilutions, the coinfection supernatants were diluted 1:10, then serially diluted 5-fold from 5^-1^ to 5^-12^. Dilutions were plated at 200μl per well of the 24-well plate and incubated at 37°C for one hour, then overlayed with 0.5 ml MEM with 3% CMC. Plaques were allowed to grow until easily visible (4-9 days). Plaques were isolated if they were a distinct, single plaque by dipping the tip of a 200μl pipet into the plaque, touching the bottom of the well, and lifting the tip out of the CMC overlay. The pipet tip was then transferred into a microtube with 200μl virus media, and gently pipetted up and down to mix the plaque isolate into the media. Each plaque was plaque isolated two additional times in the same manner, then the final plaque isolate passaged in a well of a 6-well plate to generate the virus mini stock. The mini stocks were used for viral RNA isolation for RNASeq screening for reassortment. Confirmed reassortant viruses were then selected, and full viral stocks made as described above.

ILLV and IILV were generated with our reverse genetics system in a mix-and-match transfection protocol as previously described(28). Briefly, for ILLV we transfected pINKV_L, pLACV_M, and pLACV_S, and for IILV we transfected pINKV_L, pINKV_M, and pLACV_S at 0.5μg per plasmid into BSR-T7/5 cells at ∼15% confluence in a 6-well plate. Supernatants were harvested at ≥80% CPE, and then mini stocks and full stocks of viruses grown as described above.

### Virus sequencing

For the reassortant virus screens from coinfections, viral RNA was isolated from the 6-well mini stocks using the QIAamp viral RNA mini kit following the protocol described in(30). Viruses were then screened via RNASeq to determine reassortment and purity. Sequencing libraries were generated from total RNA using the Stranded Total RNA Prep, Ligation with Ribozero Plus kit, according to the manufacturer’s protocol (Illumina). In brief, 11 μL total RNA was used as input for ribosomal depletion followed by library construction, according to the instructions. Final libraries were confirmed on Agilent’s 2100 Bioanalyzer using the High Sensitivity DNA chip (Agilent Technologies) and quantified using the Kapa Library Quantification Kit Illumina (Kapa Biosystems). Quantified libraries were combined in equimolar concentrations and sequenced as paired-end 150 bp on the Illumina MiSeq using the MiSeq Reagent Micro, V2, 300 cycle kit (Illumina). Raw reads were demultiplexed, trimmed to remove adapter sequences, and quality filtered using Cutadapt (31) and FASTX Toolkit. Reference sequences were generated for each virus by concatenating L, M, and S segment nucleotide sequences obtained from NCBI (Inkoo virus, KT288269.1, U88060.1, U47138.1; La Crosse virus, EF485033.1, EF485034.1, EF485035.1). Filtered reads were mapped to each virus reference sequence using Bowtie2 with --no-mixed --no-unal -X 1000 parameters (32). Aligned SAM files were converted to indexed BAM files using samtools (33) and coverage for each virus reference was calculated using GATK DepthOfCoverage with -omitIntervals -omitBASEOutput parameters. Coverage for each virus segment was determined using samtools idxstats (33).

Final virus stocks were sequenced by isolating viral RNA as described above. Sequence libraries were generated using 11μl of the isolated viral RNA using the Illumina Stranded Ligation, Total RNA-Seq kit with the following modifications: Ribo-Zero Plus rRNA depletion was performed, anchors were diluted 1:2 and library amplification cycles were increased to 17. NGS libraries were assessed using the Agilent Bioanalyzer 2100 and quantified using Roche’s Kapa Quant. Libraries were normalized for 2 x 150 bp cycle sequencing using MiSeq Micro v2 chemistry. Raw fastq reads were trimmed of adapter sequences using Cutadapt v. 1.12 and trimmed and filtered for quality using Fastx Toolkit v. 0.0.14. Reads were then screened against the human genome to remove host reads. Remaining reads were mapped to the reference genomes using Bowtie2 v 2.2.9. Sequence variants were called using gatk HaplotypeCaller v. 4.2.5.0 with parameter -ploidy 2 and filtered for quality and coverage using bcftools v. 1.10.2.

### Mice

All animal experiments were performed in accordance with all applicable national and institutional guidelines under Institutional Animal Care and Use protocols at Rocky Mountain Laboratories (2019-051-E and 2022-050-D) and Montana State University (2023-238-IA). For all experiments, adult C57BL/6 mice were used between the ages of 6 and 12 weeks of age at inoculation. As we have not observed a sex difference for LACV or INKV in our extensive previous studies with these viruses(2, 27, 28, 30), we used both male and female mice for these studies. For all inoculations, mice were inoculated intranasally with 1×10^4^ PFU of virus diluted in sterile 1x PBS in a total inoculation volume of 20μl under light isoflurane anesthesia. Endpoint criteria included neurological signs of disease including, but not limited to, ataxia, limb paralysis/weakness, tremors, seizures, and circling, or other endpoint criteria of moribundity or hindered mobility preventing access to food and water. All mice were humanely euthanized via isoflurane overdose and exsanguination. For neurological disease curves, mice were followed for signs of neurological disease and immediately humanely euthanized upon signs of neurological disease or other endpoint criteria. Mice that did not develop neurological disease were euthanized at 21 days post inoculation (dpi). For neurological disease curves, n=11 for LACV and LIIV, and n=12 for INKV, ILIV, LILV, LLIV, IILV, and ILLV. For time point studies, mice were euthanized at pre-determined time points spanning pre-clinical and clinical time points for viruses that caused disease. These were 6 and 9 dpi for LACV and ILLV, and 9 and 13 dpi for INKV, all other reassortant viruses, and 1x PBS mock inoculations. Mice were euthanized at their designated time point or upon showing neurological signs or other endpoint criteria regardless of dpi. For timepoint studies, n=6 designated for each virus and each time point. Due to neurological disease signs, for the LACV 9 dpi timepoint, two mice were euthanized at 7 dpi, two at 8 dpi, and two at 9 dpi, for the ILLV 9 dpi timepoint one mouse was euthanized at 8 dpi and the other five at 9 dpi, and for the ILIV 13 dpi timepoint, two mice were euthanized at 12 dpi and the remaining four at 13 dpi. All mice were euthanized, perfused transcardially with heparin-saline, and brains collected and immediately flash-frozen in liquid nitrogen. Samples were stored at −80°C until used for brain homogenate titrations.

### Replication kinetics and cytotoxicity assays

Replication kinetics were performed as previously described(2). Briefly, SH-SY5Y cells were seeded in 96-well plates at 1×10^5^ cells per well the day prior to infection. The viruses were diluted in EMEM:F-12K + 10% FBS + 1% pen-strep to an MOI of 0.01 in 50μl media which was added to each well in triplicate for each virus. Cells and virus were incubated for one hour at 37°C, washed twice, and media replenished. At 1, 6, 12, 24, 48, 72, and 96 hpi, supernatants were harvested and pooled per triplicate, clarified, and titered on Vero cells. Three independent experiments were performed.

For cytotoxicity assays, 96-well optically clear-bottomed black plates were coated with fibronectin at 16μg/ml in 100μl per well in 1x PBS and incubated at 37°C for one hour. Residual fibronectin mixture removed by dumping and wells were washed three times with 1x PBS. SH-SY5Y cells were plated at 1×10^5^ cells per well. The next day, cells were infected with the viruses at MOI=0.01 in triplicate or mock inoculated (cells + media only) controls in sextuplicate and incubated at 37°C for one hour. Virus was removed from wells, and 100μl CellTox Green (Promega) diluted 1:1000 in EMEM:F-12K + 2% FBS + 1% pen-strep was added to each well. The plates were read on a Cytation5 plate reader (Agilent BioTek) every 24 hours from 1-96 hpi. Fluorescence intensity reads were taken for each well. Fluorescence intensity in virus inoculated wells was averaged across technical replicates, then normalized by calculating fold change from the mock-inoculated cells for each virus at each time point.

### Statistical methods

All statistical analyses were conducted in GraphPad Prism v. 10.2.0. Mouse neurological disease curves were compared by Mantel-Cox survival curve analysis comparing all viruses to LACV, and all viruses to INKV. Mouse brain homogenate titers at 6 and 9 dpi were compared between LACV and ILLV via two-way ANOVA with Sidak’s multiple comparisons test. Replication kinetics assay titers were ln-transformed and analyzed via two-way ANOVA with Dunnett’s multiple comparisons test analyzing all virus titers at all time points to LACV as the reference. Replication kinetics were additionally analyzed via simple linear regression analysis of slopes on the log phase of viral replication from 6-48 hpi comparing all viruses to LACV as the reference. For all analyses, p-values <0.05 were reported as statistically significant. Cytotoxicity assays were normalized across experiments by calculating fluorescence fold change to mock-inoculated wells. Two-way ANOVA with Dunnett’s multiple comparisons test was run on the normalized data.

## Acknowledgements

We thank Dr. Karin Peterson for providing critical support for this work. We thank the animal technicians and staff at the Rocky Mountain Veterinary Branch (RML) and MSU’s Animal Resource Center (RRID:SCR_026351) which is supported by MSU’s Office of Research and Economic Development. We thank the RML Research Technologies Branch, in particular Myndi Holbrook, Paul Beare and Kent Barbian, for virus sequencing services.

## Financial Disclosure Statement

This work was supported by the Division of Intramural Research, National Institute of Allergy and Infectious Disease through AI001102 (CWW), Montana State University startup funds (ABE), and the National Institute of General Medical Sciences of the National Institutes of Health under Award Number P20GM103474 (ABE). The funders had no role in study design, data collection and analysis, or preparation of the manuscript.

## Notes

### Competing Interest Statement

The authors have declared no competing interest.

